# Colonization and transmission of the gut microbiota of the burying beetle, *Nicrophorus vespolloides*, through development

**DOI:** 10.1101/091702

**Authors:** Yin Wang, Daniel E. Rozen

## Abstract

Carrion beetles in the genus *Nicrophorus* rear their offspring on decomposing carcasses where larvae are exposed to a diverse microbiome of decomposer bacteria. Parents coat the carcass with antimicrobial secretions prior to egg hatch (defined as Pre-Hatch care) and also feed regurgitated food, and potentially bacteria, to larvae throughout development (defined as Full care). Here we partition the roles of pre- and post-hatch parental care in the transmission and persistence of culturable symbiotic bacteria to larvae. Using three treatment groups (Full-Care, Pre-Hatch care only, and No Care), we found that larvae receiving Full-Care are predominantly colonized by bacteria resident in the maternal gut, while larvae receiving No Care are colonized exclusively with bacteria from the carcass. More importantly, larvae receiving only Pre-Hatch care were also predominantly colonized by maternal bacteria; this result indicates that parental treatment of the carcass, including application of bacteria to the carcass surface, is sufficient to ensure symbiont transfer even in the absence of direct larval feeding. Later in development, we found striking evidence that pupae undergo a aposymbiotic stage, after which they are recolonized at eclosion with bacteria shed in the moulted larval cuticle and on the wall of the pupal chamber. Our results clarify the importance of pre-hatch parental care for symbiont transmission in *Nicrophorus vespilloides*, and suggest that these bacteria successfully outcompete decomposer bacteria during larval and pupal gut colonization.

**Importance:** Here we examine the origin and persistence of the culturable gut microbiota of larvae in the burying beetle *Nicrophorus vespilloides.* This insect is particularly interesting for this study because larvae are reared on decomposing vertebrate carcasses where they are exposed to high-densities of carrion-decomposing microbes. Larvae also receive extensive parental care in the form of carcass preservation and direct larval feeding. We find that parents transmit their gut bacteria to larvae both directly, through regurgitation, and indirectly via their effects on the carcass. In addition, we find that larvae become aposymbiotic during pupation, but are recolonized from bacteria shed onto the insect cuticle before adult eclosion. Our results highlight the diverse interactions between insect behavior and development on microbiota composition. They further indicate that strong competitive interactions mediate the bacterial composition of *Nicrophorus* larvae, suggesting that the bacterial communities of these insects may be highly coevolved with their host species.

## Introduction

Animals are colonized by a diverse community of bacterial symbionts that play crucial roles in their ecology and evolution [1, 2]. This has been especially well studied in insects, whose bacterial symbionts can influence traits ranging from mate and diet choice [3] to susceptibility to natural enemies [4]. Bacterial symbionts can also differ in the fidelity of their associations with their insect hosts. Endosymbionts like Buchnera in aphids, that serve obligate functions for their insect hosts by overcoming host nutritional deficiencies, are highly specific and have been aphid for millions of years [5]. At the opposite extreme, insects can retain transient associations with bacteria whose effects are more variable [6, 7, 8, 9]. Although different factors may underlie the divergent influences of bacterial symbionts on insect hosts, one key component is the way that bacteria are transmitted between insect generations [10, 11]. Whereas obligate symbionts are almost exclusively transmitted vertically, often via direct passage through eggs, more transient associations, typical of the gut microbiota, involve an external stage where bacteria are reacquired horizontally each generation via ingestion [12, 13].

Distinguishing symbionts on the basis of transmission mode (vertical versus horizontal) has been extremely useful by focusing attention on how this can align the fitness interests of symbionts and hosts [14, 15]. However, many associations between insects and their microbial symbionts fall somewhere in the middle of these strict extremes. Among diverse possibilities, trophallaxis and coprophagy occurs when bacteria are passed horizontally between individuals via oral-oral/anal contact or fecal consumption [16, 17, 18]. Similarly, horizontal symbiont transmission can take place via ingestion of the bacteria-smeared egg-coat or via consumption of bacteria-rich capsules [19, 20]. While these methods of transfer can effectively vertically transmit symbionts from parent to offspring [12], the presence of an environmental component implies that young and developing insects can be simultaneously colonized by beneficial symbionts as well as environmental bacteria that can harm the host [20, 21]. In these cases, establishment of the inherited microbiota will be dependent on the ability for inherited symbionts to competitively exclude environmental bacteria, as well as the timing and manner of their acquisition [22, 23]. Additionally, especially for holometabolous insects that undergo a complete metamorphosis, the manner of acquisition can change markedly throughout development, at one stage occurring from the mother while at later stages through alternative transmission routes [24].

Here, we examine the mechanisms of transmission and stability of the culturable gut microbiota of the carrion beetle, *Nicrophorus vespilloides*, throughout its development. This system is particularly interesting for addressing these questions given the peculiar life-history of these organisms. Nicrophorus beetles are reared on decomposing carrion where they encounter and ingest high densities of microbes [25]. Eggs are laid in the soil in close proximity to the carcass [26]. Upon hatching larvae migrate to the carcass where they both self-feed and are fed regurgitated material from the caring parents [27, 28]. Next, following an ~ 6-7 day feeding period upon the carcass, larvae cease feeding and disperse into the surrounding environment where they eventually pupate individually in underground chambers. Finally, pupae eclose into adults, and emerge from the pupal chambers to commence feeding [29, 30].

*N. vespilloides* larvae may be colonized by a varied microbiota throughout development [25, 31, 32], and this will likely be influenced by both the presence of parents and the stage of development [25, 33]. First, parents may modify the carcass microbiota by coating it in antimicrobial secretions throughout the period of parental care [21, 34, 35]. Notably, these secretions are not sterile and contain significant numbers of bacteria that can proliferate on the carcass. Secondly, parents feed larvae with regurgitated food which may facilitate the transfer of the parental gut microbiota to offspring (Post-hatch care) [36]. Finally, following dispersal, larvae cease feeding, thereby preventing continued colonization from diet-borne bacteria; and then during metamorphosis they shed the larval gut [37, 38]. At present, there is no understanding of the dynamics of these gut bacterial communities through time.

There is little knowledge of the colonization dynamics of Nicrophorus gut bacteria or the extent to which colonization is influenced by parental care, a hallmark of this system. To examine these questions we manipulated *N. vespilloides* parental care and used a culture-based approach to monitor the dynamics of symbiont colonization and stability through development. Although culturing can underestimate bacterial densities when compared to total cell counts or sequence-based approaches (see Supplemental Figure 2), this approach allowed us to examine the largest set of experimental conditions, while also identifying the bacterial groups that can be experimentally manipulated to understand mechanisms of colonization and community assembly of the microbiota using the *Nicrophorus* model system. Briefly, our results provide strong evidence that beetle parents play a defining role of the establishment of the bacteria residing in *Nicrophorus* larval guts; however, continuous parental care and feeding is not essential for the stable maintenance of this microbiota. Most strikingly, we also find that pupae undergo a sterile aposymbiotic stage, after which they are recolonized from the bacteria shed into the pupal chamber. We discuss these results in the context of the role of the *Nicrophorus* microbiota for beetle fitness.

## Methods

### General procedures

Experimental beetles were taken from an outbred laboratory population derived from wild-caught *N. vespilloides* individuals trapped in Warmond near Leiden in The Netherlands, between May and June 2014. Beetles were maintained in the laboratory at 20°C with a 15:9 hour light:dark cycle. All adults were fed fresh chicken liver twice weekly. To generate outcrossed broods, non-sibling pairs of beetles were allowed to mate for 24 hours in small plastic containers with soil. Next the mated pair were provided with a freshly thawed mouse carcass weighing 24-26 g in a 15cm x 10cm plastic box filled with approximately 1-2 cm of moist soil. Although fresh carcasses may differ in bacterial composition from aged carcasses [39], our use fresh carcasses in this study ensured higher brood success and is consistent with recent data showing that most mouse carcasses are discovered shortly after they are placed in experimental forests [40] Broods were reared in sterile soil until the point of larval dispersal from the carcass, after which larvae were transferred to new boxes for pupation with unsterilized peat soil to complete development. Soil was sterilized using two autoclave cycles at 121°C for 30 minutes, with a cooling interval between cycles.

### Maternal care manipulation

To examine the role of parental care on the acquisition and composition of beetle gut bacteria, we reared larvae under three treatment conditions that modified the degree of parental care they received [25]: 1) Full Care (FC) broods experienced complete parental care, including pre- and post-hatch care; 2) Pre-hatch parental care (PPC) broods were reared on a carcass that had been prepared by the female, after which she was removed prior to the hatch/arrival of larvae; and 3) no-care (NC) broods experienced neither pre-nor post-hatch care. Broods in all treatments were initiated similarly. Mated females were provided with a fresh carcass and induced to lay eggs. Eggs were collected and surface sterilized within 12-24 hours and these were then used to generate replicate broods of 15-20 larvae each. Females remained with their prepared carcasses in FC broods, while females were removed prior to reintroducing larvae in the PPC broods. NC larvae were provided with a freshly thawed carcass with a sterile incision in the abdomen to permit larval entry.

### Bacterial density and composition throughout development

We examined the dynamics of *N. vespiloides* intestinal microbiota through time by destructively sampling beetles throughout development. To quantify gut bacterial CFU, the whole intestinal tract from each beetle from independent broods (n = 3 at each time point) was carefully removed with fine forceps and suspended in 0.7 ml sterile sodium phosphate buffer (PBS; 100 mM; pH 7.2). The inner contents of pupa were examined in their entirety owing to the absence of a clear gut at this stage. Individual gut/pupal contents were serially diluted in PBS and plated on 1/3 strength Tryptic Soy Broth agar and incubated at 30°C. In experiments shown in Supplemental Figure 1, we also directly compared bacterial densities determined from total microscopic counts versus via plating. Although plating for CFU consistently underestimates bacterial densities, this approach recovered up to 60% of total counts and the dynamics of bacterial densities perfectly mirror those based on total counts. The composition of the maternal microbiota was characterized from n = 3 mated females.

At each time point from each treatment, we isolated random colonies (n ≥ 100) from individual beetles to analyze for species identification using MALDI-TOF Mass Spectrometry (Matrix Assisted Laser Desorption Ionization-Time of Flight) with the Biotyper platform (Bruker Daltonic GmbH). By generating unique whole-cell protein200 based fingerprints for each colony, the Biotyper permits highly reproducible identification of bacterial colonies to the genus or species level. Because of its reproducibility, ease of use and cost effectiveness, the Biotyper is used extensively in clinical and public health microbiological laboratories [41] and is finding increased use in ecological studies [42, 43, 44]. To standardize growth prior to analysis, individual colonies were tooth-picked onto a 1/3 TS plate and grown overnight. Colonies were then transferred directly to a 96-well steel MALDI-TOF target plate and coated with 1 µl of alpha-cyano-4-hydroxy cinamic acid (HCCA) matrix comprised of Acetonitrile (50%), Trifluoroacetic acid (2.5%) and water (47.5%), and dried at room temperature. The target plate was subsequently inserted into the Biotyper system for analysis. Next, mass spectrometry was carried out using the MALDI Biotyper RTC (Realtime classification) and analyzed using Biotyper 3.0 (Bruker DAltonic GmbH). Spectra were collected under the linear positive mode in the mass range of 3 to 20 kDa and a sample rate of 0.5 GS/s (laser frequency, 60 Hz; ion source 1 voltage, 20.08 kV; ion source 2 voltage, 18.6 kV; lens voltage, 7.83 kV). The Bruker bacterial test standard (BTS 8255343) was measured for standardization of MALDI calibration before the specimens were processed. Spectra were compared to the reference library provided by Bruker which identified 62.3% of the colonies to species level overall using a stringent cut-off of 1.699, below which indicated no reliable identification (in the Bruker library) [45, 46]. To confirm these assignments and to establish the identity of colonies whose spectra were not included in the Bruker database, all unique MS spectra (including both those with positive hits and those not present in the Biotyper database) were subsequently analyzed using 16s rDNA sequencing. Colony PCR using primers 27F (5’AGAGTTTGATCCTGGCTCAG-3’) and 1492R (5’-GGTTACCTTGTTACGACTT-3’) was used for bacterial 16s rRNA gene amplification [47]. The PCR cycling conditions were as follow: 95 °C for 5 min, then 34 cycles of 95°C for 45 s, 55°C for 45 s, 72°C for 1 min. PCR products were directly sequenced via the DNA Markerpoint in Leiden and 16s sequences were classified for bacterial taxonomy using a nucleotide BLAST against the NCBI database. The Bruker database was manually updated to include new samples thus obtained.

A second experiment was conducted to determine the source of bacterial re233colonization following beetle pupation. Pupae were removed from their chambers and both the inside of the chamber and the cuticle were swabbed with a sterile, moist, cotton swab. The bacteria on the swab were resuspended in sterile water and serially diluted onto 1/3 TS agar. Finally, soil from outside the pupal chamber was collected and diluted into PBS and plated. Colonies were isolated and identified as above using a combination of MALDI-TOF Biotyping and 16s rDNA sequencing. To exclude rare or transient bacterial species, we established a minimum threshold frequency of 1%, averaged over all sampling periods for each treatment set, prior to analysis of community composition.

### Statistical analysis

Bacterial CFU through time were analyzed using General Linear Models (GLM) with time and treatment as factors. Community composition was analyzed using the Vegan package in R [48]. Beta diversity among the different treatments was analyzed using ANOSIM, which is based on a Bray-Curtis dissimilarity matrix [49, 50]. The Dendrograms to examine community similarity were generated based on the matrix of mean within-group and between-group distances and the R function *hclust* was used for hierarchical clustering.

## Results

### Bacterial CFU vary through development and as a function of parental care

The CFU of intestinal bacteria was quantified throughout development for three treatment groups corresponding to different levels of parental care. Following hatching from sterile eggs, larvae from all treatments rapidly acquire high bacterial densities within their guts. Bacterial densities vary significantly through time (GLM analyses: df = 10, P < 0.001) and as a function of treatment (GLM: df = 2, P = 0.006) and vary across nearly 6 orders of magnitude as a function of developmental stage. These dynamics are insensitive to experimental methods, as estimates of density based on total microscopic counts perfectly mirror those determined by plate counting (Supplemental Figure 2). During larval feeding on the carcass, bacterial densities increase in all treatments, reaching densities of ~ 10^6^ / larva. By contrast, following dispersal, bacterial populations precipitously decline until, during pupation, bacteria were undetectable. Finally, as pupae eclose and reemerge from pupal chambers, they reacquire a high-density bacterial population within their guts (Fig. 1). It is notable that this recovery occurs prior to feeding and before emergence from the pupal chamber, indicating that recolonization takes place from bacteria resident within the pupal chamber itself.

**Figure 1.**
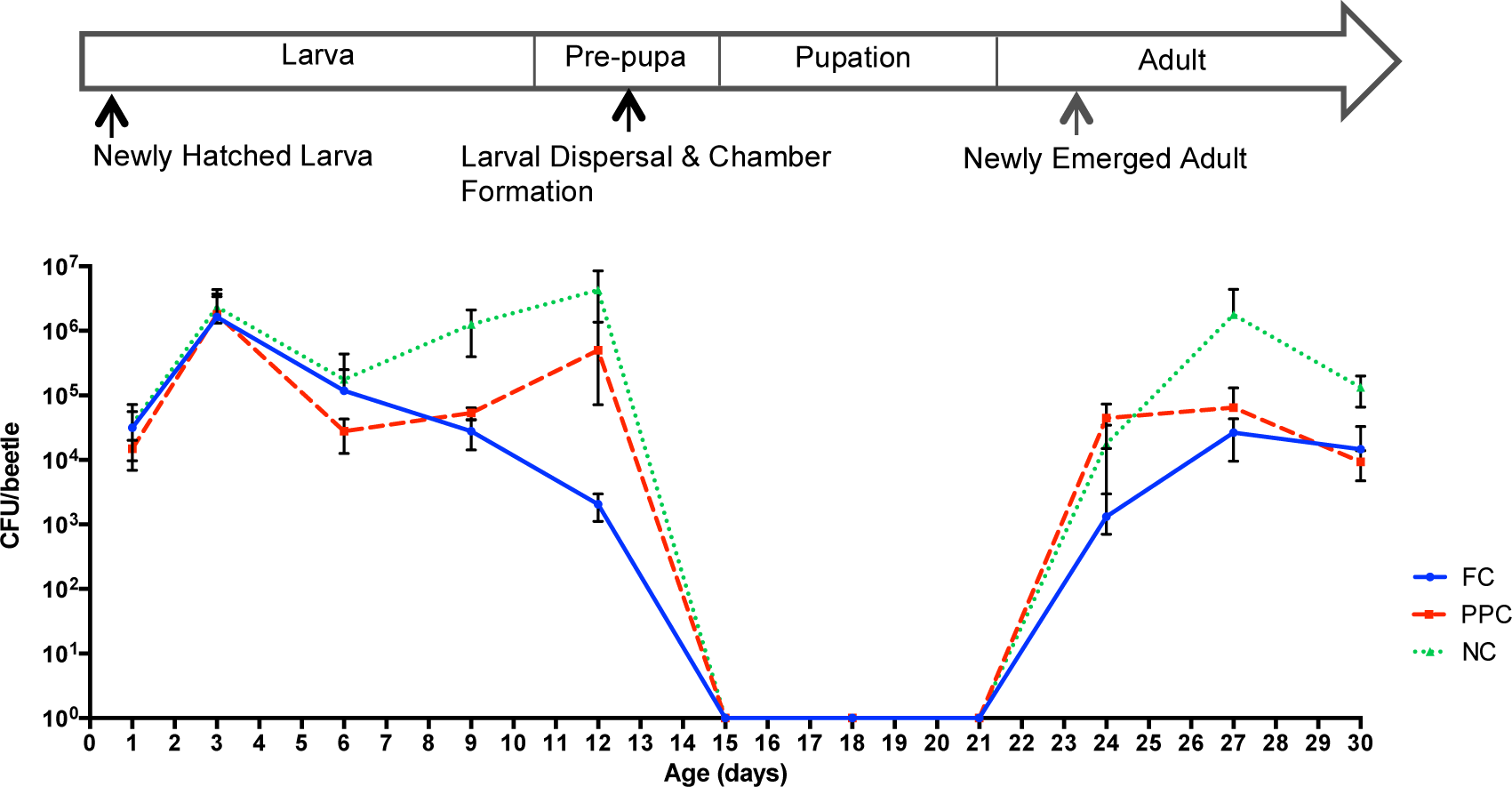
CFU of *Nicrophorus vespilloides* gut bacteria through development. Overview of the entire time course of beetle developmental and change in CFU of host gut contents through time (means ± SD). FC corresponds to larval guts sample from full parental care broods; PPC, to guts sample from pre-parental care broods; and NC from larval guts sample from no-care broods.

### Composition of *N. vespilloides* larval symbionts

Although bacterial densities differ across parental-care treatments there is broad overlap in the dynamics of CFU change through time. Despite these similarities, the composition and diversity (Table S1) of these communities may vary. To understand these differences and to illuminate transmission dynamics from mothers to larvae, we tracked community composition of gut bacteria within larvae throughout development (Fig. 2) using MALDI-TOF Mass-Spectrometry and compared these to the maternal samples. The maternal microbiota was dominated by four bacterial Genera that together comprised > 65% of recovered CFU, including *Providencia, Morganella, Vagococcus* and *Proteus*, with several other genera appearing in lower frequencies (Fig. 2). We next examined genus level composition across the three larval treatment groups. As anticipated if transmission occurs via parents, we observed significant overlap in the bacterial communities of parental and larval gut communities from larvae receiving parental care throughout development (R_FC-g vs Mother_=0.277; P=0.028, Table 1), as R values < 0.25 correspond to “barely separable” groups [51]. Equally, although to a lesser degree there is concordance between the maternal microbiota and those of larvae receiving pre-hatch care only (R _PPC-g vs Mother_= 0.331; P = 0.066, Table 1). By contrast, larvae reared in the absence of parental care are highly diverged from the parental microbiota (R _NC-g vs Mother_= 1; P = 0.007, Table 1) (Fig. 3A and 3B). In particular, the gut community of NC larvae was shifted towards bacterial groups likely acquired from either the soil or the carcass (Fig. 2 and 3C), e.g. *Escherichia coli* (23.5%), *Serratia* (20.4%) and *Staphylococcus* (19.2%).

**Figure 2.**
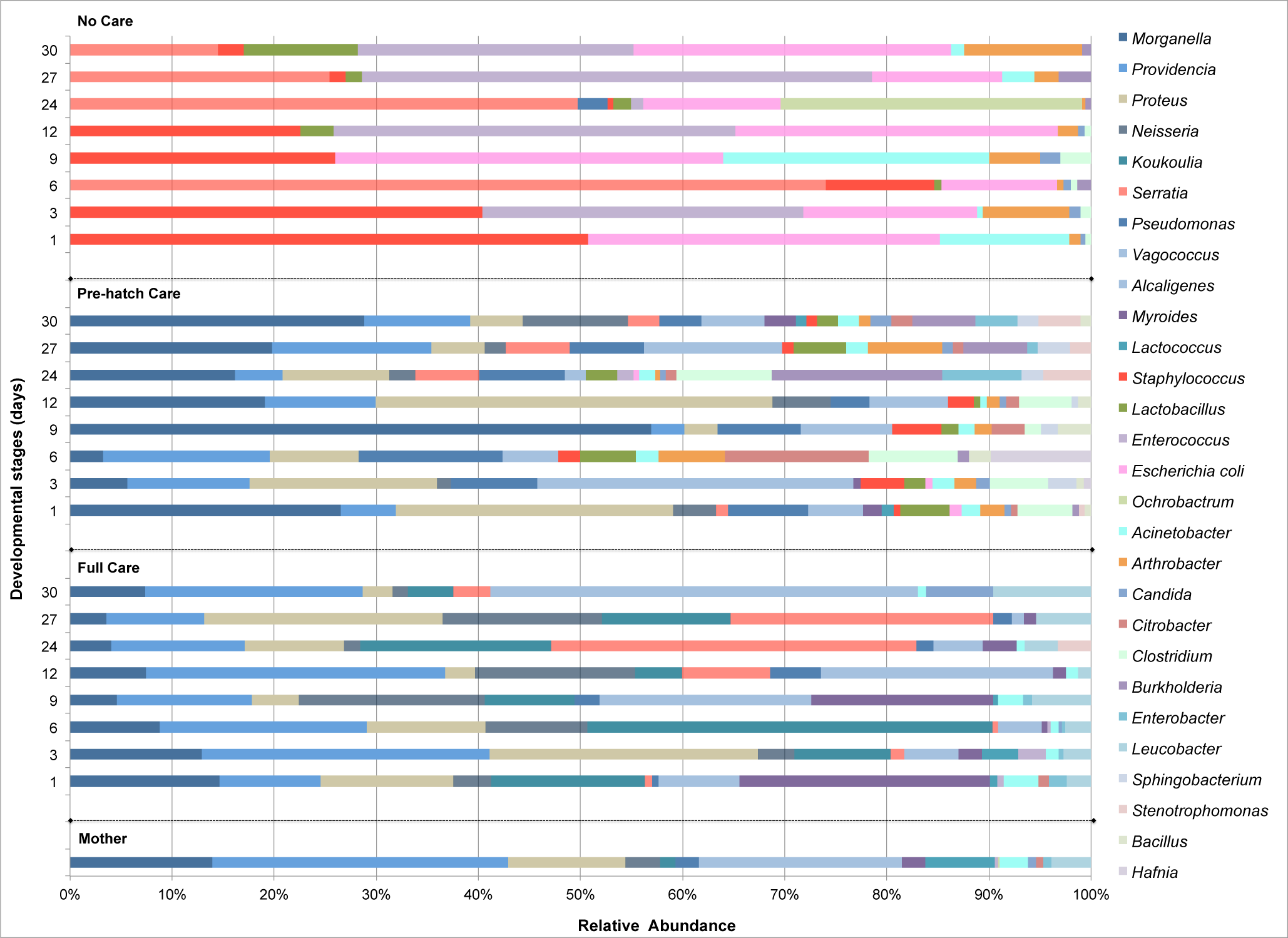
Composition of *N. vespilloides* gut microbiota through development. The maternal gut microbiota is shown at bottom, while treatment designations are the same as in Figure 1. No CFUs were detectable between days 15-21 of larval development, corresponding to the duration of pupation. Three individual larvae were independently analyzed for each time point. Y-axis of day 1 to day 9 refers to larval stage; day 12 corresponds to prepupal stages; day 24 to day 30 refers to adult formation, respectively.

**Table 1.** ANOSIM analysis on bacterial community dissimilarity. Subscripts correspond to the site of isolation; e.g. FC-g corresponds to gut samples, FC-c corresponds to the pupal carapace and the wall of the pupal chamber, and Soil corresponds to bulk soil outside the pupal chamber.

| Groups | R statistic | Significance (P value) | Number of permutations | Number of observed | Test model |
| --- | --- | --- | --- | --- | --- |
| FC-g, PPC-g, NC-g | 0.8152 | 0.001 | 999 | 352 | Global |
| FC-g, PPC-g, NC-g, Mother | 0.741 | 0.001 | 999 | 264 | Global |
| FC-c, PPC-c, NC-c, Soil | 0.7493 | 0.001 | 999 | 814 | Global |
| FC-g, NC-g | 0.9556 | 0.001 | 999 | 144 | Pairwise |
| PPC-g, NC-g | 0.9939 | 0.001 | 999 | 64 | Pairwise |
| FC-g, PPC-g | 0.6651 | 0.001 | 999 | 144 | Pairwise |
| FC-g, Mother | 0.2769 | 0.028 | 999 | 24 | Pairwise |
| PPC-g, Mother | 0.3306 | 0.066 | 999 | 24 | Pairwise |
| NC-g, Mother | 1 | 0.007 | 999 | 24 | Pairwise |
| FC-g, FC-c | 0.3177 | 0.068 | 999 | 54 | Pairwise |
| PPC-g, PPC-c | 0.0242 | 0.052 | 999 | 54 | Pairwise |
| NC-g, NC-c | 0.0134 | 0.397 | 999 | 54 | Pairwise |
| FC-c, Soil | 0.8889 | 0.094 | 720 | 9 | Pairwise |
| PPC-c, Soil | 1 | 0.114 | 720 | 9 | Pairwise |
| NC-c, Soil | 1 | 0.099 | 720 | 9 | Pairwise |

In comparing the larval microbiota of the three treatment groups, ANOSIM analysis illustrated clear differences between the treatment groups overall (Global test: R = 0.815, P = 0.001) and although there are differences between the FC and PPC larvae, there is much greater similarity between the two groups with parental care (R_Fc-g vs ppc-g_ = 0.665; P = 0.001) compared to either care group and the no-care larvae (R_FC-g vs NC-g_ = 0.956, P=0.001; R_PPC-g vs NC-g_ = 0.994, P=0.001) (Table 1, Fig. S1). This is also apparent in the Venn diagrams in Fig. 3A, focusing on presence/absence of specific bacterial groups. Together, these results indicate that transmission of the beetle microbiota occurs predominantly from parents to offspring. However, they also reveal that continued replenishment of bacteria from parent to offspring via feeding is unnecessary to establish the endogenous microbiota. Instead transmission can occur indirectly via deposition of the maternal bacteria on to the carcass by the mother during carcass preparation and subsequent colonization of larva via self-feeding.

**Figure 3:**
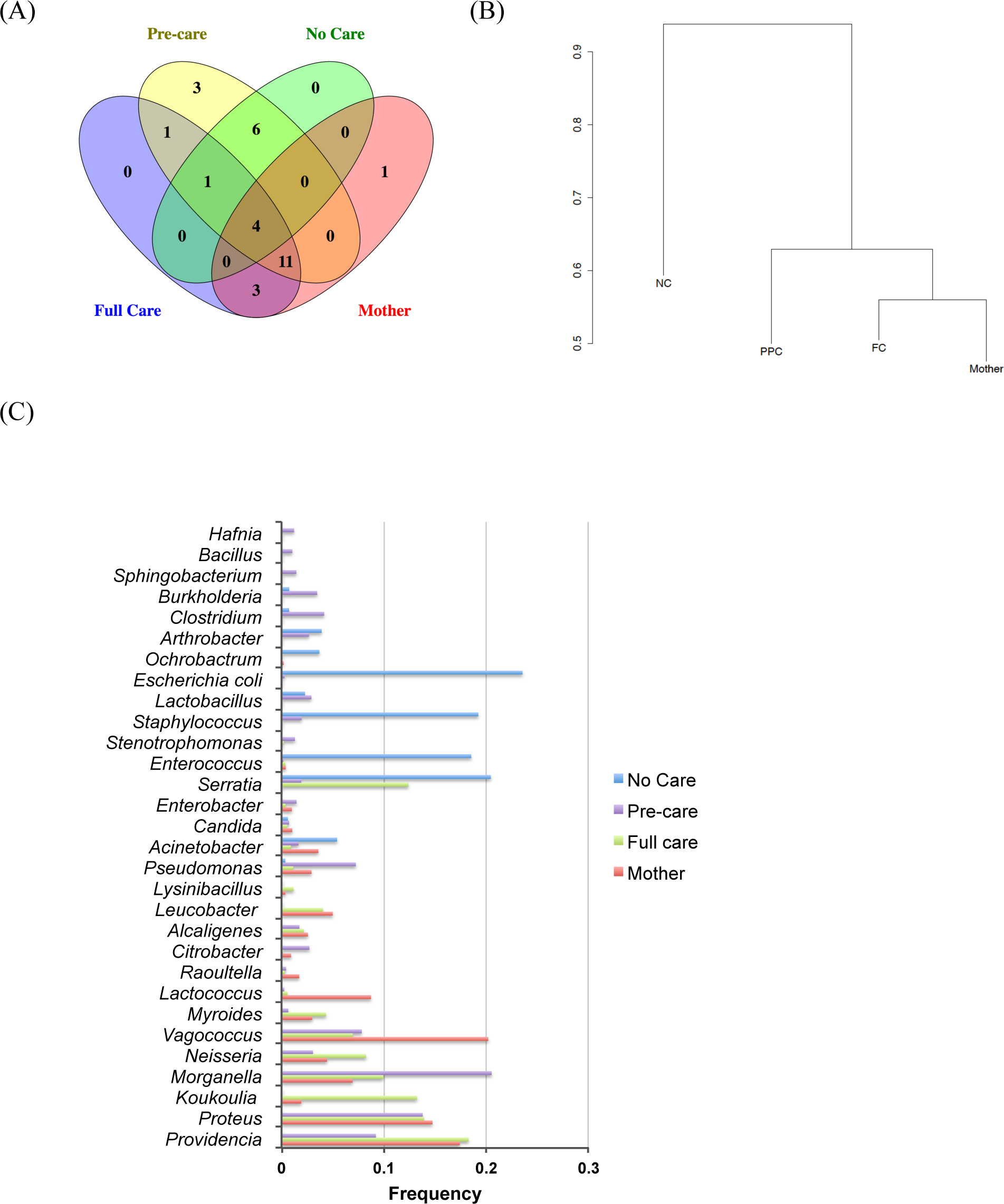
Frequencies of bacteria from gut communities across parental care treatments. **A)** Shared and unique genera between treatment groups. Strains with a minimum frequency of 1% were included. B) Hierarchical clustering on mean similarity of gut microbiota between treatment groups. **C)** Overall composition of gut communities across treatments. Strains with frequencies lower than 1% across all communities were excluded from plots.

### Re-colonization of *N.vespilloides* symbionts

A striking result from these analyses is the aposymbiotic stage occurring during pupation, followed by recolonization from within the pupal chamber. Notably, this result based on CFU was further confirmed by direct microscopic counts (Supplemental Figure 2). To assess the source of recolonization, we sampled bacterial populations from the pupal cuticle and the wall of the pupal chamber, together with samples from the bulk soil in which pupal chambers were constructed. Treatment designations are as above, with the addition of subscripts corresponding to each sampling site. For example, FC-g refers to samples taken from the guts of larvae receiving Full Care, while FC-c represents samples from the cuticle and chamber wall of these same larvae. These analyses showed that the *N. vespilloides* pupal cuticle and chamber soil had very similar compositions (FC-g, FC-c: R = 0.32, P = 0.068; PPC-g, PPC-c: R = 0.02, P = 0.052; NC-g, NC-c: R = 0.03, P= 0.0397 by Pairwise test of ANOSIM, Table 1), and that these were diverged compared to the bulk soil (FC-c, Soil: R = 0.89, P = 0.094; PPC-c, Soil: R = 1, P = 0.114; NC-c, Soil: R = 1, P = 0.099 by Pairwise test of ANOSIM, Table 1). Importantly, many bacterial genera irrespective of treatment, were found in the pre-pupal gut and the cuticle but infrequently or not at all in the soil. For example, the most common bacterial groups in FC larvae contained *Providencia* (FC-g: 18.3% vs FC-c: 17.1%), *Morganella* (FC-g: 10.0% vs FC-c: 8.7%), *Proteus* (FC-g: 14.0% vs FC-c: 3.9%), *Vagococcus* (FC-g: 7.0% vs FC-c: 6.1%), *Neisseria* (FC-g: 8.3% vs FC-c: 4.7%), and *Koukoulia* (FC-g: 13.2% vs FC-c: 8.9%), while these were absent from soil. Similarly, the most abundant genera in NC beetles were only found in NC-g and NC-c: *Escherichia coli* (NC-g: 23.5% vs NC-c: 23.8%), *Enterococcus* (NC-g: 18.5% vs NC-c: 18.8%) (Fig. 4A and 4B). These results indicate that the core components of previously colonized gut bacteria can successfully recolonize the host intestinal system after the aposymbiotic stage characteristic of pupation. Thus, although transmission and recolonization to larvae occurs via the environment, the bacterial species that recolonize the newly eclosing adult are highly biased towards bacterial species that were already present in the pre-pupal gut and which were originally acquired from the mother.

**Figure 4:**
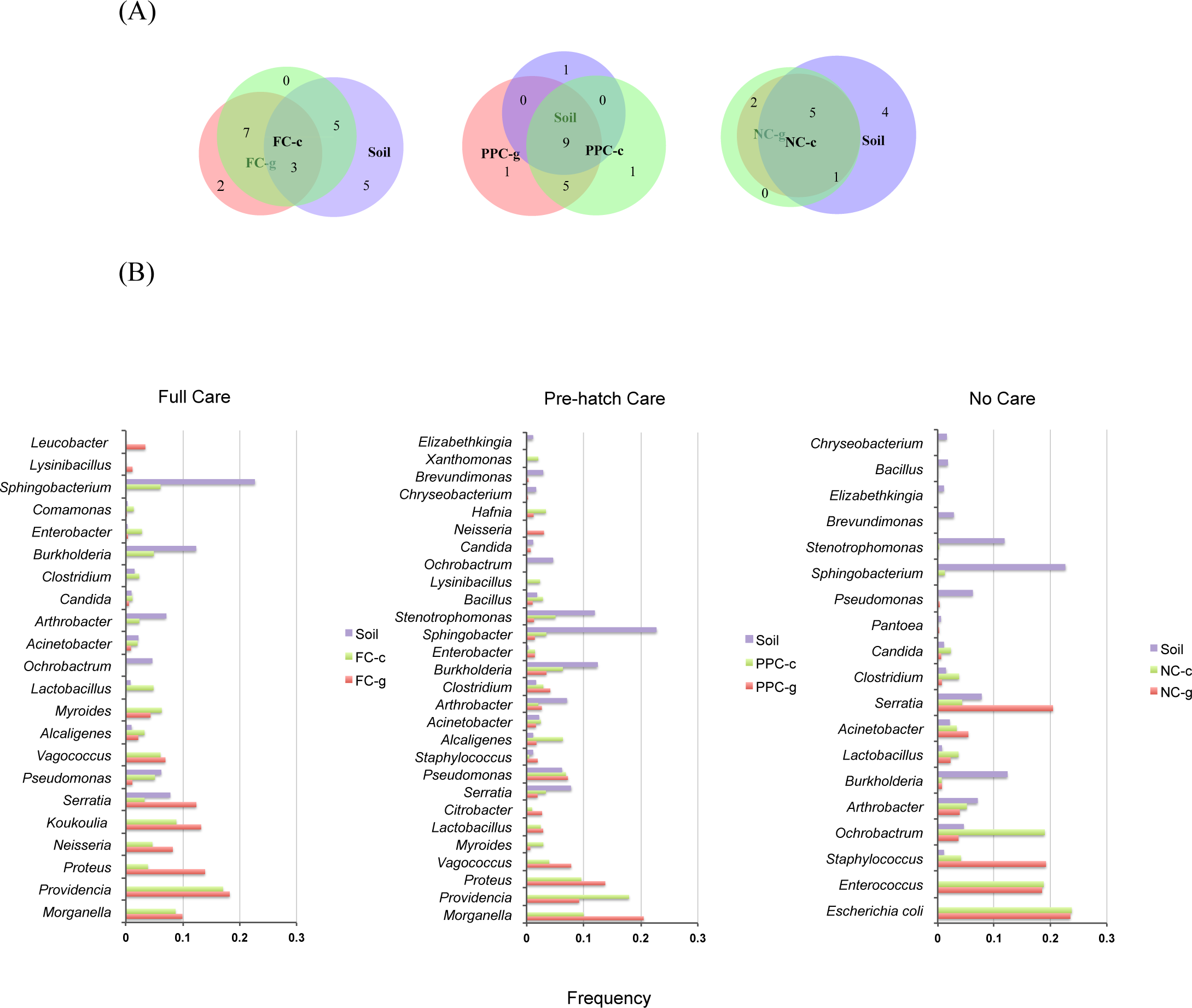
Re-colonization of bacterial communities through pupations. A) Shared and unique genera among treatment groups. Subscripts correspond to the site of isolation; e.g. FC-g corresponds to gut samples, FC-c corresponds to the pupal carapace in the wall of the pupal chamber, and Soil corresponds to bulk soil outside the pupal chamber. **B)** Comparison of gut bacterial communities from each sample site and treatment. Strains with frequencies lower than 1% across all communities were excluded from plots.

## Discussion

Animal symbionts can be passed to offspring through different mechanisms that vary in their reliability of transmission [10, 11]. While strict endosymbionts of animals are typically transmitted vertically via eggs, other mechanisms that include an environmental component may also reliably transmit bacteria between generations [14, 19]. Here, we examined the mechanisms of bacterial transmission from *Nicrophorus vespilloides* mothers to offspring. *Nicrophorus* larvae are exposed to and consume high densities of bacteria throughout their development on decomposing carrion [25]. In earlier studies we showed that parental care, including preservation of the carcass through secretion of lysozyme and potentially other antimicrobials, is essential for maintaining larval fitness [21]. Additionally, preliminary metagenomic analyses from our own lab (unpublished data) and published studies from others [39] have found that parental beetles significantly modify the bacterial composition of decomposing carrion, thereby potentially influencing the bacteria that larvae are exposed to and ingest.

To examine the influence of parental care on the transmission of bacteria from parents to offspring, we manipulated the level of care parents provide to their larvae. With full parental care, parents apply oral and anal secretions to the carcass both before larvae hatch and throughout larval development [21, 32]; they also regurgitate food to larvae during the first three to four days of development [27, 52]. As expected, given the continuous direct and indirect exposure to parental bacteria, larvae in this treatment were colonized predominantly with parental symbionts (Fig 2); importantly, despite limitations associated with a CFU-based approach, we observed broad overlap between the dominant bacterial species we cultured and those identified using sequence-based approaches (e.g *Providencia, Morganella, Vagococcus, Proteus, Koukoulia*, and *Serratia)* [53]. However, with this Full Care treatment alone, it could not be determined if larvae require constant replenishment of the parental species for these to be maintained in the larval gut [54]. One possibility, for example, is that the dominant bacteria from the carcass could outcompete endogenous beetle bacteria within the larval gut; this could be driven actively, if the bacteria on the carcass are particularly good colonizers, or passively since larval exposure to carcass bacteria is continuous [22, 23]. To address this question, we established broods that only received pre-hatch care. In this treatment, parents have no direct exposure to larvae, and can only influence larval exposure to bacteria indirectly through their influence on the carcass. It is important to note that because eggs are sterile, transmission is also prevented through this route [26]. As with the Full Care treatment, larvae receiving only Pre-hatch care were also predominantly colonized by maternal bacteria (Fig. 2 and 3). This was not due to an inability of bacteria from the gut to colonize larvae, as larvae in the No-Care treatment were also colonized by a high-density bacterial microbiota. Also, bacteria in the pre-hatch groups were partially colonized by carcass-derived bacteria (Fig. 2 and 3C), leading to higher bacterial diversity overall in this group (Supplemental Table 1) and indicating the capacity for carcass-derived bacteria to establish themselves within the larval gut. Rather, we interpret this result to indicate that “endogenous” bacteria from the mother are able to outcompete the carrion associated microbes. Furthermore, this effect is long-lasting and can persist entirely in the absence of direct maternal feeding. Although this interpretation is consistent with our data, this hypothesis will require experimental testing using the culturable species we have now established in our collection of *N. vespilloides* symbionts.

At present, we understand relatively few of the mechanisms used by parents to manipulate the carcass bacteria. However, several factors are likely to be important. First, when parents locate a carcass they strip it of fur, while simultaneously coating the carcass surface with oral and anal secretions. The composition of these secretions has only been partially characterized, but a key component is lysozyme, a broad-spectrum antibacterial with greater specificity towards Gram-positive bacteria [32, 55]. Additionally, oral secretions contain bacteria that can serve as an inoculum to feeding larvae (unpublished results). In addition to these behaviors, we have also observed parents opening the carcass and removing the mouse gut, behaviors that could potentially have a dramatic influence on larval bacterial exposure by introducing oxygen that could bias the bacterial community towards aerobic species or more simply by directly reducing the overall density of bacteria to which larvae are exposed. Following gut removal, parents continue to coat the carcass in secretions and then bury the balled up carrion underground [21, 29, 52], which could influence moisture or temperature levels. Both behaviors could possibly bias the persisting microbial species, and potentially in favor of species originally introduced by caring parents. In addition, caring parents and their larvae may be exposed to different bacterial numbers and composition as a function of carcass age, a factor that is known to have a dramatic influence on larval fitness [25, 56]. Although much remains to be determined of these processes, our results clarify the importance of more completely understanding how parents influence both the bacteria on the carcass and how this, in turn, affects larval microbiota establishment.

After larvae complete feeding, they migrate into the soil to pupate [29, 52]. Bacterial numbers during this stage decline precipitously (Fig. 1 and Supplemental Figure 2), in part due to the absence of feeding and also to the evacuation of the larval gut. In addition, larvae in some metamorphosing insects undergo a pre-pupal molt which would further reduce bacterial numbers [50, 57]. Regardless of the mechanisms, it appears that *Nicrophorus* larvae become effectively sterile during pupation, an outcome previously seen in several flies and mosquitoes [58, 59, 60]. It is possible that host immunity facilitates pupal symbiont suppression during metamorphosis [57, 61, 62], as a decline of phagocytic haemocytes and an increasing phenoloxidase activity were both detected in *Nicrophorus* pupa [63]. Following this aposymbiotic state, bacterial densities are quickly recovered at eclosion with bacterial communities that significantly overlap with those present prior to pupation (Fig. 1 and 2). To determine the source of recolonization, we sampled bacteria from the pupal molt as well as the wall of the pupal chambers, and in both cases we observed striking similarity to the microbiomes of earlier developmental stages. Interestingly, this was true for all treatment groups, suggesting that there is no intrinsic bias to recolonization, but rather that eclosing beetles are colonized by a subset of the bacterial species present in the pupal chamber.

The larval gut of *N. vespilloides* thus appears to be colonized via a combination of mechanisms that are dependent on the degree of parental care and the stage of development. With complete parental care, parents transmit bacteria to larvae through a combination of direct feeding and through an indirect effect mediated by the carcass [36, 39]. At present, it remains unclear if this latter component is because *Nicrophorus* symbionts outcompete the mouse carrion microbiota within the larval gut, or if this occurs primarily on the carcass surface itself. However, the former seems more likely given the vast differences in larval exposure to these two groups of bacteria, and the fact that larvae in the pre-hatch group remained colonized by beetle symbionts, despite lacking any direct exposure to parents (Fig. 2). It is tempting, given the reliable mode of transmission from parents to larvae, to speculate about the function of these sybmionts for *Nicrophorus* growth and development, particularly the role of these bacteria in limiting infection from carrion-borne bacteria [39, 53]. However, this remains an active area of research that we will hope to address in future publications. In addition, it will be important to supplement the present work with more detailed analyses based upon sequencing [39, 53]. Although culture-based methods play an essential role in unraveling the relationships between invertebrate host sociality and their symbiont strain-level diversity [64], they are clearly complementary to sequence-based methods that can recover bacterial groups that may be difficult or impossible to culture in the laboratory. Our work clarifies the key links between *Nicrophorus* social behavior and symbiont transmission. This is likely to have parallels in other animal systems where parents invest in the care of offspring.

## Acknowledgements

Funding for this work was provided by start-up funds from Leiden University to DR and from a studentship from the China Scholarship Council to YW. We gratefully acknowledge expert assistance with beetle maintenance from Kees Koops, and thank Andres Arce and Chris Jacobs for their extremely helpful comments on an earlier version of this manuscript. In addition, we gratefully acknowledge the helpful and constructive comments from the Editor, Eric Stabb, and three anonymous reviewers on an earlier submission of this manuscript.

